# SCHEPHERD: A universal platform for high-throughput, high-resolution, and programmable control of cell behavior through bioelectric stimulation

**DOI:** 10.1101/2025.09.16.674226

**Authors:** Yubin Lin, Jeremy S. Yodh, Celeste Rodriguez, Paul Kreymborg, Daniel J. Cohen

**Author notes:** These authors contributed equally (Yubin Lin, Jeremy S. Yodh). Correspondance (YL), (DJC).

## Abstract

Spanning frogs, fish, and humans, direct-current (DC) bioelectric cues play critical roles beyond neuro-muscular function, such as modulating morphogenesis, immune response, and healing through electrotaxis—electrically directed cell migration. Harnessing this potential requires new tools. However, standardized, accessible, and reproducible infrastructure capable of DC stimulation remains a challenge. We present SCHEPHERD: a universal, electrobioreactor integrating 8 stimulation channels and modular inserts to enable most electrotaxis assays in one device (cells, monolayers, and 3D spheroids), while enabling powerful, new capabilities. SCHEPHERD revealed through parameter sweeps that DC fields act like a ‘steering wheel and gas pedal’ for cell migration. We then used live confocal imaging to observe electrically reprogrammed F-actin dynamics. Finally, our multi-polar inserts generated complex spatial electrical patterns that reorganize engineered tissue dynamics. By significantly improving accessibility through modularity and an open-source, graphically programmed stimulator, we hope SCHEPHERD can help broaden the community studying these important DC bioelectric phenomena.

## Introduction

Bioelectricity plays a critical role beyond neuro-muscular processes^1^, including programming cell migration^2–4^, regulating healing and disease^2,5–10^, and directing morphogenetic processes^5,11–14^. Here, the electrical cues are distinct from familiar, charge-balanced action potentials—rather they consist of slow-changing, direct current (DC) ion gradients producing fields of ∼ 1 V/cm *in vivo*^1,2,9,12^. These DC cues act on membrane-bound proteins to trigger *electrotaxis*—the directed migration of cells down an electric potential gradient—in nearly every tested model system, spanning slime molds^15^, zebrafish^5^, frogs^4,12^, and mammalian models^2,10,13,16–18^. The myriad roles of electrotaxis have sparked significant interest in better stimulation tools to study the mechanisms of electrotaxis^2,4,16,18–21^, control engineered tissues and organoids^3,13,16,17,22^, alter development *in vivo*^5,12^ and accelerate healing^2,6,7,10,23,24^.

Unfortunately, while existing neuromuscular stimulation technology is safe for the rapid pulsing needs of action potentials, the same hardware causes toxic redox reactions when used for sustained DC stimulation^25,26^, necessitating development of bespoke devices that have been difficult to reproduce, control, scale, and share. As a result, there is currently no ‘standard’ platform for electrotaxis, limiting reproducibility, trust, and progress^6,27,28^. These significant barriers-to-entry have greatly hindered progress in this exciting field.

Here, we present a new approach for high-throughput, reproducible, and computer-programmable DC stimulation for cells, tissues, and organoids in multiwell plates. This platform—SCHEPHERD (Spatiotemporal Cell Herding with Electrical Potentials for High-throughput Electrobiology with Reproducible Design) —not only performs nearly all previously published live-imaging electrotaxis protocols in an integrated platform, but improves on them through simpler set-up and new capabilities not possible with other approaches. These capabilities include: high-throughput parameter sweeps that reveal hidden behaviors; live-imaging of real-time directional control of the cytoskeleton; and production of precise, spatially patterned electric fields that drive local behaviors within tissues similar to large-scale optogenetics.

Our goal with SCHEPHERD is to provide a foundational platform to comprehensively serve, and broaden, the growing field of bioelectricity and to accelerate near-term work on wound healing, immune modulation, and tumor suppression. To enable this, SCHEPHERD is not only entirely open source hardware/electronics, but addresses a key accessibility concern for biology labs: we are committed to providing and supporting our custom electrical control boards and 3D-printed infrastructure, which should allow a new level of standardization, reproducibility, ease-of-setup, and capabilities not possible in other systems.

## Results

### SCHEPHERD is an integrated platform that solves key problems in the field and enables new capabilities

What fundamentally sets DC bioelectricity apart from the more familiar AC (neuromuscular action potentials) is that it is a slow, steady process driven by sustained ion currents. However, slow and steady currents in an electrolyte will cause toxic redox reactions if not carefully managed, so all DC bioelectric systems follow a strategy similar to our approach in Fig. 1. First, a power supply drives current through non-redox electrodes (e.g. Ag/AgCl^29–31^, conductive polymer^32–37^, etc.). These electrodes drive continuous ion flows carried through saline buffers and hydrogel salt bridges both couple to the living sample and filter out any harmful byproducts. Current must then be concentrated over the biological sample as it is current density (current/cross-section) that drives many DC biological processes^24,27,38–41^, so samples are typically grown in microfluidic environments with small cross-sections. A detailed discussion of design parameters has previously been presented^3^, but the key design guideline is shown below in Equation 1.

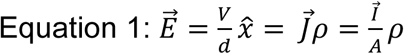

**Figure 1:**
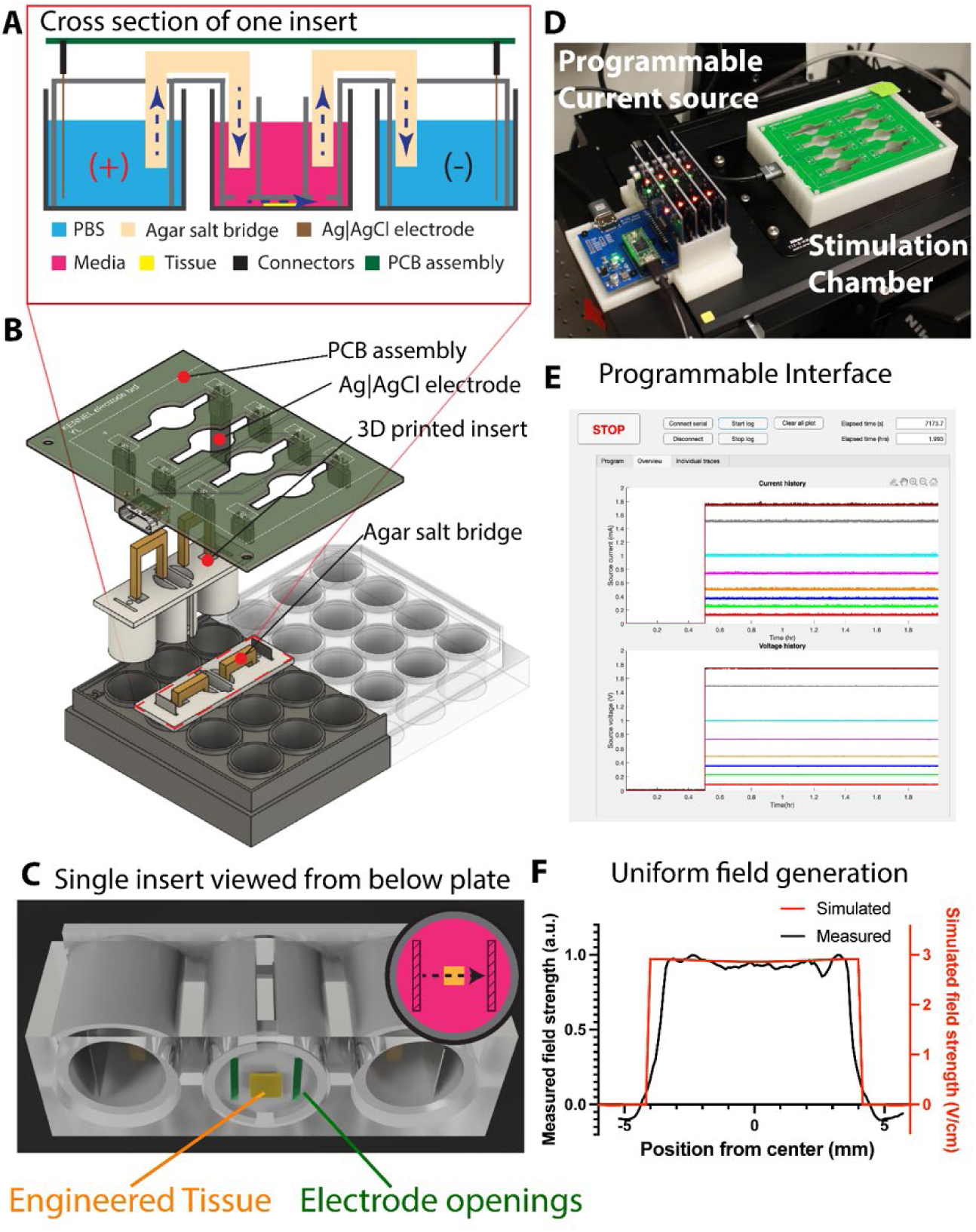
Device and platform overview. A. Cross section schematic of a single insert. A voltage drop is applied across an Ag (Anode) and Ag/AgCl (Cathode) electrode pair generating an ionic current from PBS wells (blue), through agar salt bridges, into a media well (pink) containing a monolayer tissue (yellow). B. 3D rendering of the multi-well insert. 3D printed inserts are slotted into a standard 24-well plate, agar bridges connect the PBS wells to the media well, and a PCB lid plate which contains the Ag and AgCl electrodes is placed on top of the 24-well plate. C. 3D rendering of the bottom of a 3D printed insert showing how the field flows over a tissue. The insert has a lip around its perimeter used to maintain a ∼250 μm gap for current to flow through the two current openings. D. SCHEPHERD on the microscope showing the enclosed 24-well plate with PCB lid plate assembly and the programmable current source. E. Screenshot of the custom software used to control the 8 independent stimulation. The specific current and voltage traces were used for the electric field sweep in Fig. 3. F. The electric field in the x-direction from a COMSOL simulation (red) shows homogenous stimulation across the well, which we validated with charge beads (black).

Where: 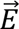= electric field strength (V/cm); *V* = voltage drop across the channel; *d* = length of culture channel, 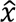 = unit vector in the x-direction; 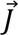 = current density; *I* = current (mA); *A =* cross-sectional area of the channel (mm^2^); and ρ = resistivity of the culture media (Ωm). Typical field strengths for DC cell biological experiments target O (1 V/cm). See prior work for more details^3,16,18^.

Our approach is to encapsulate the entire stimulation process in a modular, 3D printed format that can be slotted into a variety of multi-well plates (adjusted accordingly) and essentially dropped directly on top of cultured cells, tissues, organoids, etc. that are already growing in the multi-well plate. Figs. 1A-C demonstrate how our approach can be implemented within a 24-well plate format. Here, the stimulation module occupies three contiguous wells within the plate, with the outer two wells containing paired Ag/AgCl electrodes immersed PBS (brown and blue, resp. in Fig. 1A), and the center well contains the sample to be stimulated immersed in cell culture media (yellow, Fig. 1A). We cast two agarose salt bridges in acrylic plastic molds so that they perfectly slot into, and bridge the electrode wells to the culture well in the middle, providing a continuous electrical circuit and maintaining electrochemical separation (beige, Fig. 1A). Our monolithic, 3D printed insert is designed to settle into the plate under its own weight and naturally form the ceiling and walls of a precise tunnel with a height of 250 μm (adjustable) around the cultured sample (yellow, Fig. 1A). Openings in the ceiling of this central insert allow ionic currents to flow through this effective microfluidic channel and to stimulate the sample with high current density. Fig. 1B shows how multiple modules can slot into a multi-well plate, while Fig. 1C shows a view from below a mounted insert where a square cultured tissue is stimulated between two parallel electrode openings in the ceiling. The direct integration into multi-well plates allows us to leverage existing, standardized culture systems and support most forms of inverted, live-cell imaging. The field uniformity is verified with electrophoresis from the charged fluorescent beads (Fig. 1F), and the result agrees with the computer simulation, see methods for details.

Up to 8 of these three-well modules can be loaded simultaneously in a 24-well plate (see Fig. 1D), which we made far more manageable by introducing a custom printed-circuit-board (PCB) ‘lid’ that incorporates all the external wiring, automatically aligns the Ag/AgCl electrode films in each well, and provides space for salt bridges and transmitted imaging (green PCB in Fig. 1B, 1D). This minimal lid also allows the platform to more easily integrate into live microscopy set-ups, and the concept can be generalized for a variety of multi-well plate formats. All modules are fully programmed, monitored, and driven through a single cable that connects the PCB lid directly to our custom power supply.

The final element of any DC electrobioreactor is the power supply, which ultimately limits the kind of stimulation that is possible. In practice, the majority of DC stimulation studies rely on repurposed laboratory power supplies^38,42–47^ (often lacking proper safeguards and resolution) or professional current sources^17,24,40,48^ (costing thousands of dollars). While custom solutions exist^41,49^, no extant system (including our own prior work) effectively combines high-throughput, programmable, safe, and fully-logged DC stimulation as the hardware does not exist. Moreover, the wide variation in power supply setups across studies can make it difficult to replicate prior studies or even set up a safe stimulation system. We solved this issue by designing and integrating a custom, 8-channel current source with independent programming and data logging for each channel (Fig. 1E, S1, and S2). This system was built from the ground up for DC stimulation during live microscopy. Power delivery is handled through precision current sources for constant current control, which is far more reproducible for DC electrobiological use and also compensates for the interfacial resistances between electrodes and electrolytes to guarantee a stable electric field across the sample throughout an experiment. The miniature format of our inserts allows for low operating voltages (up to 20 V), thereby improving safety by reducing shock hazards^47,50^, and our single-cable connection between this custom power supply and the stimulation plate eliminates the wire and salt bridge tangle typical of prior electrotaxis systems (including our own). Now, a 1 mA driving current will produce ∼2 V/cm within a stimulation well, and this can be independently adjusted for up to 8 wells. This entire interface is a fraction the cost of commercial solutions while enabling unique functions, and a complete bill-of-materials and design guides are provided in our supplement. Fig. 1D illustrates the complete system setup that includes the stimulation controller, the stage top live imaging SCHEPHERD, and a surrounding enclosure for CO2 delivery and temperature control.

### Validation of the DC stimulation for common experimental approaches

We next validated an individual stimulation module (the 3-well insert) against challenging cases from the literature involving unidirectional electrical stimulation that would normally require different set-ups.

We first targeted generating a uniform electric field across an engineered tissue by designing an insert with parallel electrode openings in the ceiling to produce a uniform electric field (Fig. 2A). We tested against large-scale collective electrotaxis^3,13,16,51^ as this is relevant for a range of studies involving healing and development (Movie S1). Here, a whole tissue is stimulated with a uniform electric field to drive collective cell migration from anode to cathode. Collective electrotaxis is an important validation case both because of the scale of the sample (e.g. cm) and the need for the electric field to be uniform over that large area. To demonstrate this, we engineered a precise epithelial monolayer of primary mouse keratinocytes 1.5×6 mm in size (see methods) and drove its collective electrotaxis with a 3 V/cm field (Fig. 2B). As shown in Fig. 2C, we observed tissue-scale collective migration, demonstrating both the expected increase in speed with electrotaxis and highly directional motion (aligned parallel to the applied electric field). SCHEPHERD mainly uses fluorescence imaging to maximize its flexibility. However, with minimal modification, the concept of SCHPHERD was easily adapted for transmitted-light imaging. Fig. S5 shows the schematic of the insert, where two pieces of acrylic provide imaging window for the transmitted light path. We tested this with MDCK cells under 3 V/cm stimulation in Movie S2.

**Figure 2:**
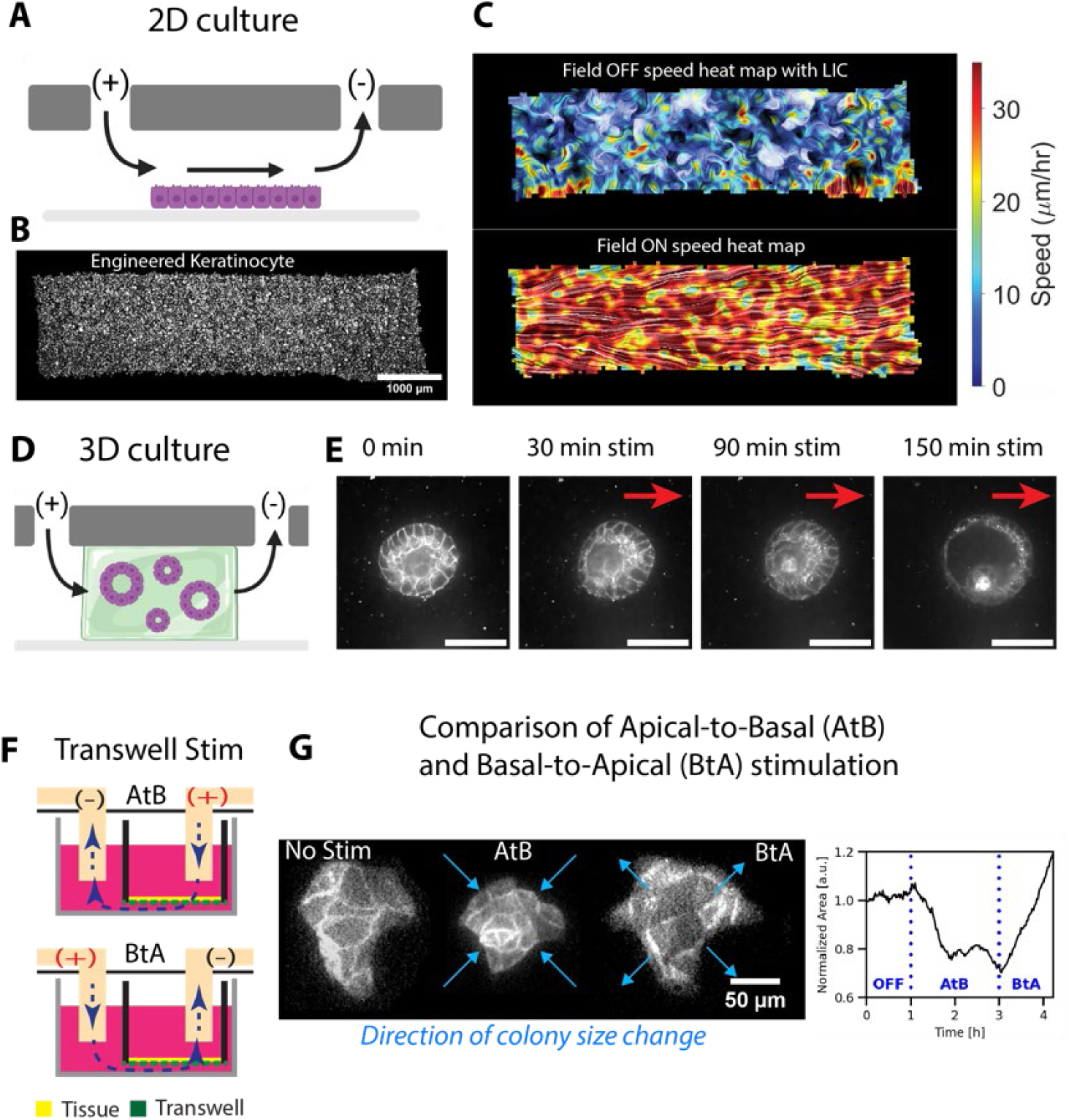
Reproducing the canon of electrotaxis results with the SCHEPERD assembly. A. Schematic of homogenous current flow over a tissue monolayer. B. A micrograph of a 6 mm x 1.5 mm monolayer tissue of primary mouse keratinocytes (see Methods). Scalebar is 1000 um. C. Migration flow pattern (see Methods) overlaid on a speed heat map without electric stimulation (top panel) and with electric stimulation (bottom panel) shows a uniform response to the field. Color corresponds to speed magnitude and texture shows streamlines. D. Schematic of electrically stimulating 3D kidney cysts embedded in a hydrogel (see Methods). E. Confocal slice over time of electro-inflation of the cyst. Scalebar is 50 um. F. Schematic of transverse (vertically oriented) stimulation through a transwell showing Apical-to-Basal (AtB) and Basal-to-Apical (BtA) stimulation. G. Micrographs of an MDCK colony which contracts with apical-to-basal (AtB) stimulation and then expands with basal-to-apical (BtA) stimulation. The normalized area dynamics are plotted showing AtB induces contraction and BtA induces expansion of the island.

Our second test case was 3D DC stimulation of ‘epithelioids’—hollow, cyst-like structures of epithelial cells grown in hydrogels that have previously been shown to both inflate (‘electroinflation’) and break symmetry during DC stimulation^13^. This type of stimulation was previously performed with confocal imaging and continuous media perfusion in a relatively complex device (our former design). To accommodate the working distance required for confocal imaging, we grew the tissues in a hydrogel within the well of a glass-bottomed 24-well plate, then then gently slotted the stimulation insert into the well to form the stimulation channel around the tissues (Fig. 2D). Subsequent electrical stimulation recapitulated the previous data, showing rapid inflation and symmetry breaking (thinning on the anode side, thickening on the cathode side; Fig. 2E, Movie S3).

Finally, we tested transwell stimulation—an advanced culture mode where a semi-permeable culture membrane separates upper and lower media chambers that is used for electrical stimulation in complex culture environments^52,53^ and for transmigration experiments with immune cells^54,55^. Here, we modified our printed insert to accommodate a commercial transwell platform (see Methods) and allow us to direct an electric field perpendicular to cultured cells (Fig. 2F). Previous work with epithelial stimulation has shown that stimulating in the Apical-to-Basal (AtB; top-to-bottom) direction causes cells to contract, while stimulating Basal-to-Apical (BtA; bottom-to-top) causes cells to expand^56^. Our data replicates this demonstrating strong contraction and expansion of islands of epithelial cells over hours of AtB/BtA cycles (Fig. 2G, Movie S4, S5).

Together, these three demonstrations show how SCHEPHERD can recapitulate prior results from a wide range of different culture and stimulation conditions all within a single platform, emphasizing that one platform can handle the majority of different experimental needs.

### SCHEPHERD enables high-throughput DC bioelectric experiments

We next evaluated new capabilities enabled by SCHEPHERD. A key challenge in DC bioelectricity is relating the stimulation input for a given model system to specific output (e.g. migration speed) given the wide range of possible electrical inputs. While resistive ladder strategies have been demonstrated in the past for single cells^41,46^, high-throughput stimulation of engineered tissues is much more difficult and prior work in this space typically sticks to previously established field strengths^3,16,18,51^. Here, we tested and demonstrated the high-throughput capabilities of SCHEPHERD to sweep 7 different field strengths from 0.25 V/cm-3 V/cm in a single experiment using our multi-channel, 24-well plate from Fig. 1. The final three wells were used as negative controls (0 V/cm). Here, the plate was configured as shown in Fig. 3A, and to further emphasize throughput, we engineered 3 replicate tissues per stimulation well (Fig. 3B, Movie S6). This allowed us to capture a complete sweep of tissue responses within a single experiment with multiple replicates per condition. Figs. 3C, D show the dynamics of cell migration speed or directionality (0 = undirected motion, 1 = fully aligned with the electric field vector). The high throughput of this sweep allowed us to construct a ‘transfer function’ plot (Fig. 3E) cleanly mapping the directionality and speed of the bulk of the tissue to a given electrical input. It must also be noted that performing this kind of sweep with any extant system would have taken at least one month of microscope time and sample prep to serially run the experiments and replicates, whereas we were able to complete all replicate sweeps in a single day of back-to-back imaging.

**Figure 3:**
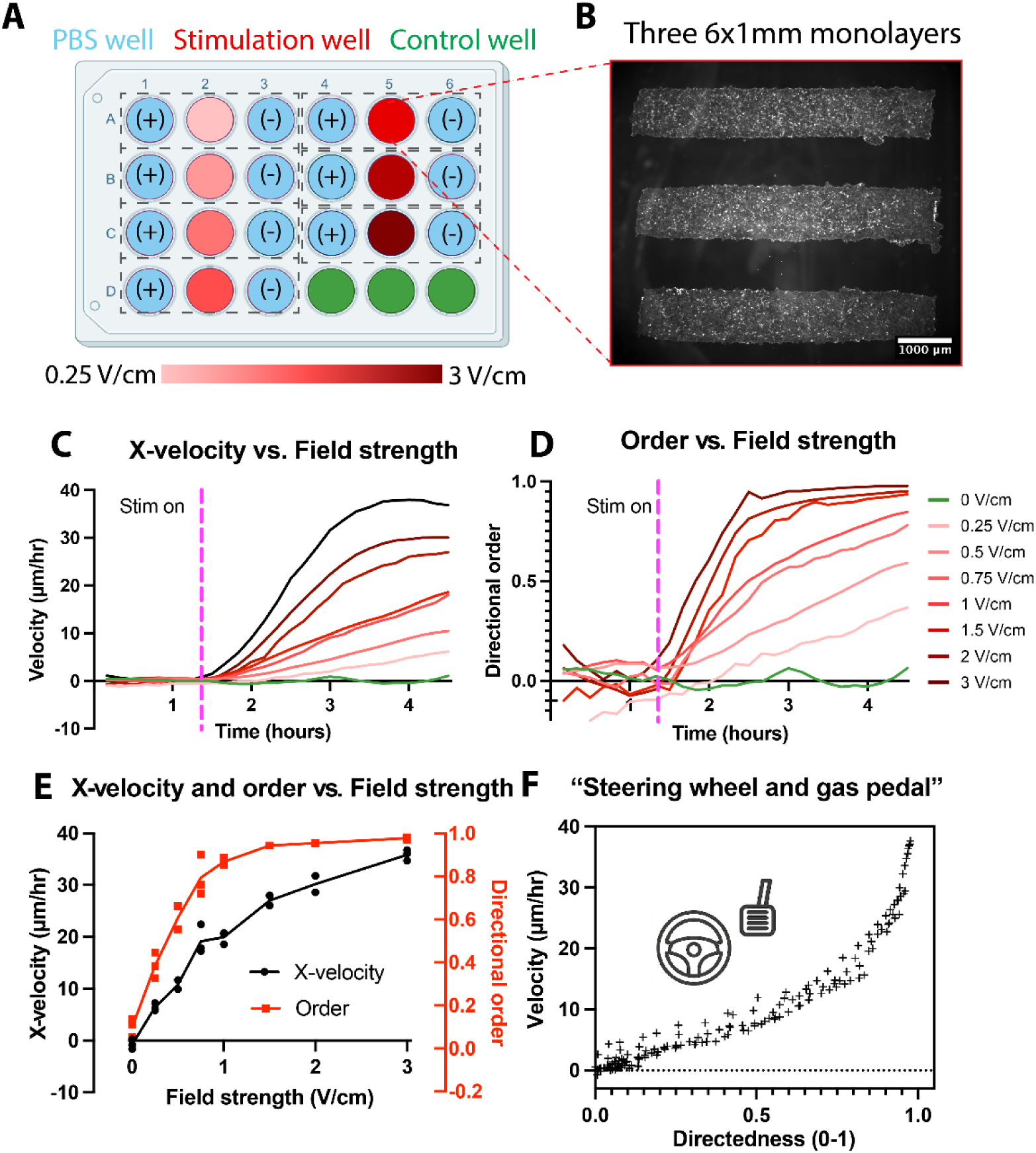
SCHEPERD device massively increases experimental throughput. A. 24-well schematic of an electric field amplitude sweep. Each triplet of wells contains an independent stimulation module (as in Fig. 1A) with an anode (+) and cathode (-) pair flanking a stimulation well. With this assembly, a 24-well plate contains up to 8 stimulation triplets, allowing for 8 independent experimental conditions; here, three wells were devoted to negative controls. In the schematic, color corresponds to field strength. B. A micrograph of one well with three 6 mm x 1 mm MDCK tissues. Scale bar is 1000 μm. C. The dynamics of the x-component of the velocity for the stimulation sweep show that increased field strength results in increased x-velocity. D. Dynamics of the directional order (measured with respect to the applied electric field) as a function of field strength that the directional order saturates faster with stronger fields. E. Transfer function of the x-component of the velocity and the directional order as a function of electric field strength. Data points were taken three hours into stimulation. F. Speed plotted as a function of directedness showing at least two distinct regimes of behavior. For relatively low directional order (<∼0.8), speed increases linearly with directional order. At higher values of directional order, (>0.8), the speed scales super-linearly with directional order. These data emphasize our ‘steering wheel and gas pedal’ framing of electrotaxis.

However, these finely mapped data raised another question—why do cells significantly speed up during electrotaxis, as shown in our transfer function (Fig. 3F) and as noted in prior studies^3,18,51^? One hypothesis in the field is that electrical stimulation causes Joule heating of the media around the cells and that migration scales with temperature^4^. We ruled that out here as part of our validation of SCHEPHERD where we measured temperature in the tissue culture channel during stimulation and observed no obvious temperature shifts (Fig. S4).

A competing hypothesis, known as the ‘compass model’ of electrotaxis^51,57^, is that speed increases *because* directedness is increasing. Essentially, as the direction of motion of individuals in a crowd becomes more uniform, there should be less frequent neighbor-neighbor collisions, which increases movement efficiency and allows cells to reach higher speeds. If true, this hypothesis suggests that speed should saturate when directedness saturates, i.e., when directedness reaches ∼1, speed should plateau. A direct way to test this argument is to correlate speed with directionality for all datapoints from our field sweep (see Fig. 3F), where each data point is described by a speed and a directedness. Here, we see that speed and directedness are linearly related for low values of directedness (<0.8); but as directedness reaches its maximum of 0.8-1, speed dramatically increases with a larger characteristic slope (though we note that this behavior is nonlinear). This finding proves that the ‘compass model’ of electrotaxis is incomplete. We instead suggest thinking of tissue-scale electrotaxis as a ‘steering wheel and gas pedal’ process. The direction of the electric field sets the migration direction (steering wheel), and the magnitude of the electric field sets speed (gas pedal). At low fields (*E* < 1 V/cm), migration speed increases from both the steering wheel and gas pedal, but at high fields (*E* > 1 V/cm), when steering can no longer boost efficiency, migration speed still grows via the gas pedal.

### SCHEPHERD enables complex cytoskeletal control

One of the more surprising elements of electrotaxis is how quickly and precisely it can reprogram cell migration at large scales in response to programmed electric fields. For instance, it has been previously been demonstrated that electrotaxis can not only drive monolayer of tissues across a Petri dish^3,6,18^, but can drive those sample tissue sheets to make 90-degree turns, or even spin in complete circles if the field direction can be programmed^16^. This could be a powerful approach for studying cytoskeletal remodeling as well as programming migration during morphogenesis or healing, but the previous system, SCHEEPDOG, was difficult to implement and did not support high resolution fluorescence imaging^16^. Here, we leveraged the drop-in nature of the SCHEPHERD inserts and advanced power supply enable a new, much more accessible way to do real-time cell migration control while also supporting high-resolution fluorescence imaging.

We implemented full 2D electrical control simply by adding a second axis of electrodes to our SCHEPHERD insert (see Fig. 4A) that we can independently control, meaning we can set any arbitrary electric field direction by shifting the relative weights of how strongly biased the electrodes are and which are ‘+’ or ‘-‘ as shown in the cartoon in Fig. 4A,B. Moreover, the assembly of this new insert is functionally identical to the simple, uni-axial case in Figs. 1-3 (albeit with a lower throughput since more wells are used by a single insert). To ensure this approach worked, we first demonstrated programming a ‘triangular’ maneuver (Fig. 4B) into an array of cultured primary mouse skin cells and plotted the average of their tracked trajectories over an 8 hour period (Fig. 4C; see Methods, Movie S7).

**Figure 4:**
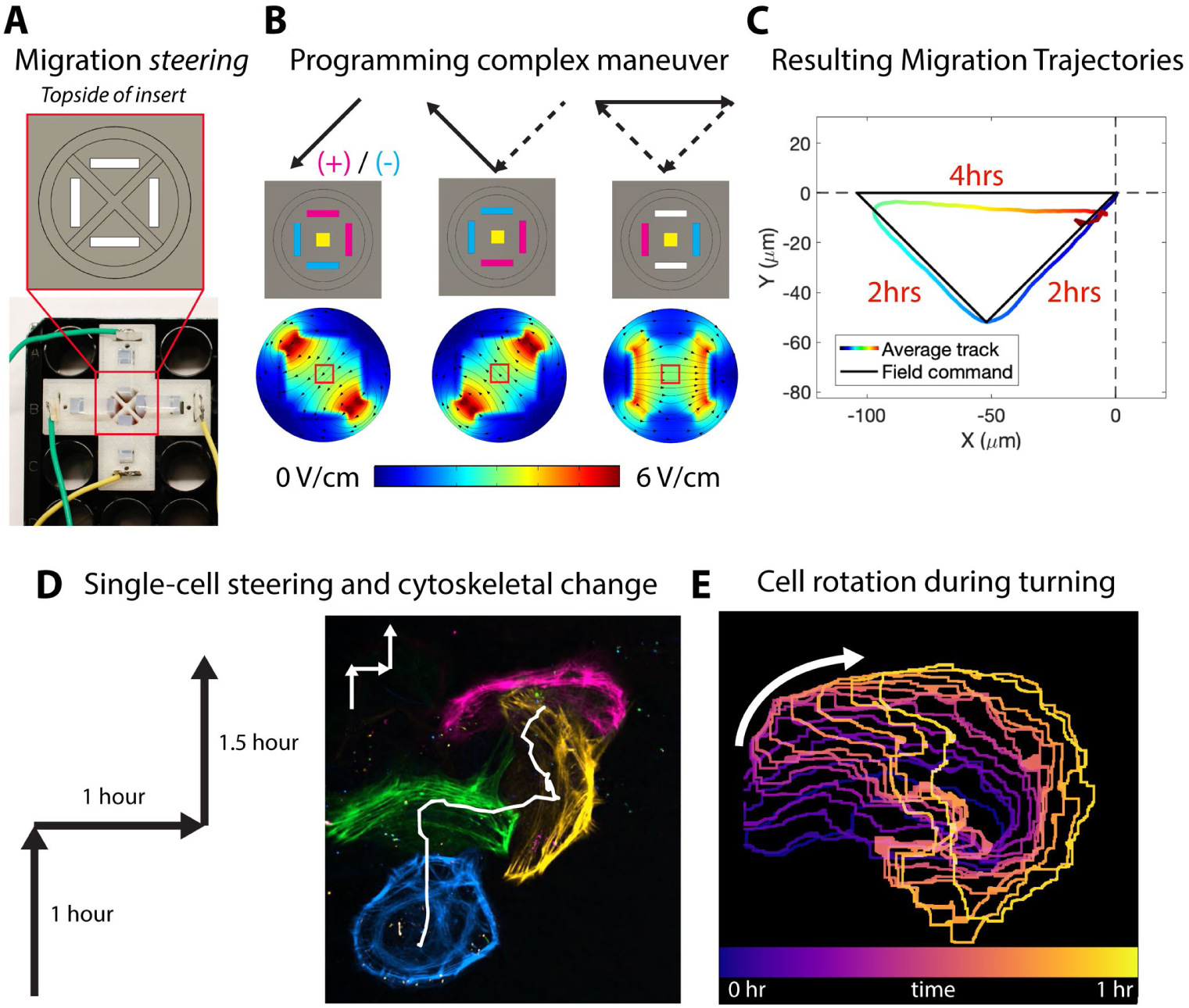
Electrically steering single cell migration and cytoskeletal dynamics in 2D. A. Schematic of the top of insert (top) and the physical experimental set-up (bottom) for XY electric stimulation. B. Top row: programmed command vectors. Middle row: anode and cathode addressing schematics. Bottom row: 2D COMSOL simulations showing the field strength and direction. C. Trajectories averaged over a population of primary mouse keratinocytes in time subject to the 2D triangular migration command from Fig. 4B. D. Left: programmed zig-zag command for single cell electrotaxis. Right: High magnification imaging of the cytoskeleton where color corresponds to time. The white solid line shows the trajectory of the centroid of the cell. E. Cell rotating during a right turn color corresponding to time.

Such extreme migratory plasticity must ultimately come from cytoskeletal remodeling, and while fixed cell studies have hinted at various mechanisms, it has been difficult to observe cytoskeletal remodeling live during electrical stimulation. Our 2D programming approach was well-suited here since we could use a ‘right’ command to first polarize cells rightward and then switch the field ‘up’ and watch how the lamellipodial machinery at the front of the cell reconfigures. This benefits from high-resolution fluorescence imaging, so we performed this experiment using a glass-bottomed 24-well plate with live epifluorescence imaging of F-actin in primary mouse skin cells (see Methods) and a programmed ‘zig-zag’ maneuver (see Fig. 4D; Movie S8). Interestingly, we observed the lamellipodium ‘swinging around’ the cell to accommodate the 90-degree turns rather than retracting and repolarizing. Fig. 4E highlights this whole cell rotation demonstrating how the lamellipodium gradually rotates clockwise in response to the ‘right turn’ field vector.

### Locally directing cell migration inside living tissues

*In vivo* data demonstrate that naturally occurring DC fields tend to have spatiotemporal patterns, as might be expected from the complex shapes, sizes, and functions of living tissues^5,12,58^. However, nearly all electrotaxis studies have focused on global stimulation with uniform electric fields (including everything shown so far in this work), in large part due to experimental constraints. SCHEPHERD’s ceiling-based electrode system offers a totally new approach to locally shaping the electric field. SCHEPHERD offers two independent ways to locally control electric field geometries based on the unique way that the ceiling works in the 3D printed insert.

First, we can directly control the number, size, shape, and location of the individual electrode apertures, all of which can be simulated computationally to design specific behaviors into a tissue. To demonstrate this, we created an insert with a ceiling containing 4 independent, small electrode openings aligned over small regions of a much larger tissue (Fig. 5A). Our simulation indicated that activating these electrodes in a unique grouping of paired anodes and cathodes (Fig. 5A) could induce complex local migration dynamics within a 5×5 mm tissue sheet of primary skin cells, and our experimental data validated this. Fig. 5B and Movie S9 reveal a complex pattern of converging and diverging cell migratory flows within the tissue, with the rainbow colors indicating the local migration axes orientations. Beyond simply controlling direction and speed, these complex patterns have implications for local cell density (Fig. 5D) and even overall tissue shape, as the patterned fields ultimately sculpt the global shape and size of the tissue.

**Figure 5:**
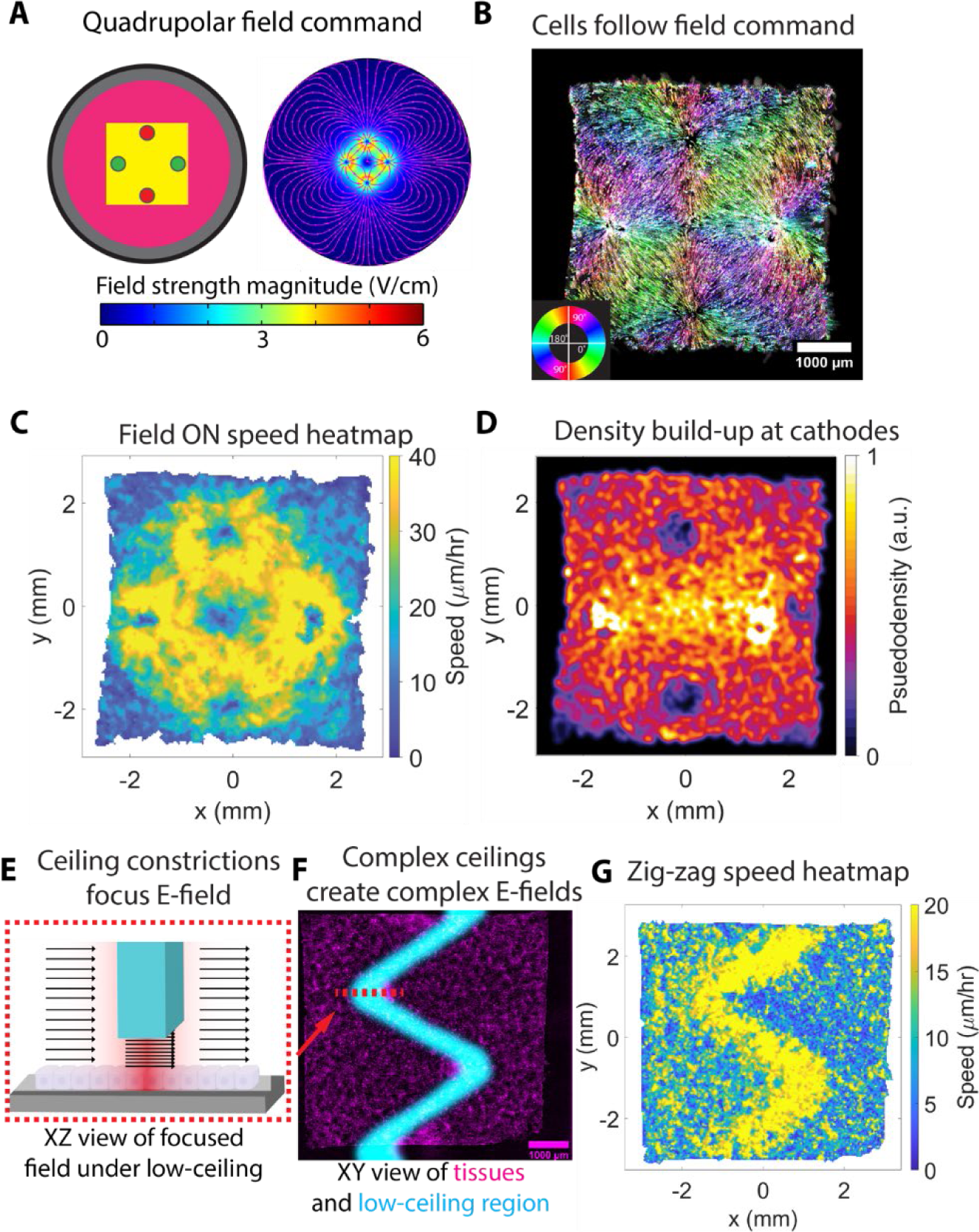
Spatial electric fields generate complex, designable migration patterns inside living tissues. A. Schematic of the anode and cathode geometry in the insert ceiling to generate a quadrupolar field (left) and a 2D COMSOL simulation of the field magnitude and direction for the field (right). B. Flows and orientations of cell trajectories show that the cells follow the field lines produced by the insert. Color corresponds to orientation. Scale bar is 1000 μm. C. Speed heatmap taken an hour into stimulation showing quadrupolar migration structure. D. Plot of the pseudo-density of nuclei shows build-up in cell density at the cathodes and decreased density at anodes. Color corresponds to pseudodensity. E. Cross-section schematic demonstrating how changing the ceiling topography can locally amplify electric field strength wherever the ceiling is closer to the floor. Image shows how the current density (arrows/region) is low everywhere except under a narrow 3d printed region allowing for local electrotaxis. F. A micrograph of a 6×6 mm MDCK monolayer subject to a zig-zag stimulation region (blue region in the micrograph). Scale bar is 1000 μm. G. Accompanying speed heatmap showing that migratory speed increases are localized to the zig-zag region. Color corresponds to speed.

Controlling the 3D topography of the ceiling offers a second approach to even more precisely and locally control tissue dynamics by creating intense, precise concentrations of current density only specific locations with far greater precision than the first method. To demonstrate this approach, we lowered a portion of the ceiling in the pattern of a ‘zig-zag’ (Fig. 5E, 5F), filmed the resulting tissue dynamics, and then produced a heatmap indicating the steady state collective migration speed pattern that resulted within the larger tissue (Fig. 5G; Movie S10).

## Discussion

Bioelectricity beyond the brain is implicated in so many critical processes— immune activity^11,59,60^, wound healing^6,23,58^, development^61,62^, transplant viability^63,64^, and cancer treatment^65–68^—that new, purpose-built tools are needed to fully capture the potential and push the field forward. While SCHEPHERD is not perfect, we hope it is a positive step and solid foundation towards advancing the field both by making these studies significantly more accessible to a broader, interdisciplinary community, and by enabling genuinely new capabilities that expand the envelope of what we can do with bioelectric stimuli.

That said, what are some remaining limitations of the SCHEPHERD platform and how might they be addressed? The most obvious is that we have currently only tested SCHEPHERD using a cage incubator built around a timelapse microscope to maintain physiological temperatures as this is the instrumentation we had available and cage-style incubation does not restrict how much instrumentation or space a sample or apparatus can occupy. However, the modularity of SCHEPHERD is such that it should be readily adaptable to small-footprint, stage-top incubators for live imaging through the use of 3D printed adapters. Additionally, there is still a small amount of electronics assembly needed. In terms of the stimulation itself, SCHEPHERD’s programmable power supply was deliberately built to favor the slow, DC fields measured *in vivo* rather than AC stimulation that is more common for neuromuscular stimulation or tumor-treating-fields to disrupt mitosis. However, the SCHEPHERD culture system itself can be directly integrated with any commercial power supply, such as high frequency AC units if needed. Perhaps the most frustrating part of SCHEPHERD reflects a challenge in the whole field of DC bioelectricity—how do we safely generate DC fields without toxic redox? While the Ag/AgCl electrodes used in SCHEPHERD remain the “gold-standard” for biocompatible stimulation^29–31^, this wasteful of silver and the total charge that can be delivered is limited as the AgCl layer is sacrificial and eventually depletes. However, exciting work is taking place to develop next-generation biocompatible electrodes for DC stimulation through the use of conductive polymers and pseudocapacitors^32–37^. As these come online, they can be directly swapped into the SCHEPHERD architecture.

Overall, we are genuinely excited to release and share SCHEPHERD as it currently represents the most reliable, user-friendly, versatile, and high-throughput platform we have tested and will hopefully enable far more researchers to enter this field. It supports from single cell to 3D tissue models, allows up to 8 simultaneous stimulation patterns, and enables real-time, shaped control of the electric field. Moreover, we are committed to making SCHEPHERD accessible and supporting its further development both through direct interactions and by sharing all the necessary plans, schematics, circuitry, and software to run SCHEPHERD and start herding cells.

## Supporting information

Video S1: Uniform uniaxial stimulation

Video S2: Uniaxial stimulation with transmitted light imaging

Video S3: 3D Cyst electrotaxis and electro-inflation

Video S4: Colony edge dynamics under paracellular stimulation

Video S5: Large MDCK colony under paracellular stimulation

Video S6: Amplitude sweep with MDCK replicates

Video S7: Biaxial tissue-level simulation

Video S8: Cytoskeletal rearrangement under bi-axial stimulation

Video S9: Complex dynamics and tissue sculpting in quadrupolar fields

Video S10: Localized electrotaxis using modified ceiling

## Acknowledgments

Support for this work was provided in part by Eric and Wendy Schmidt Transformative Technology Fund, NIH Award R35 GM133574-07, and NSF CAREER Award 2412942. We also thank members of Cohen Lab for advice and support.

## Data availability

Key microscopy data will be available on Zenodo (DOI pending).

## Code availability

All control codes and schematics for the current source, and CAD files necessary to perform the experiment will be uploaded to GITHUB (link pending).

## Methods

### Device manufacture

Each stimulation insert was 3D printed using white PLA filament (Polylite PLA natural, Polymaker) using a 3D printer equipped with 0.4mm nozzle (X1C, Bambu Lab) and with a slicer setting of 80% infill, 0.08mm layer height, and support. A PETG filament (Polylite PETG grey, Polymaker) was used for interfacing the support and the main structure for ease of removal. After printing, the supports were removed using a tweezer and then each insert was dialyzed in 500ml of deionized (DI) water overnight. Incubator enclosure was also 3D printed with a white PLA filament (Polylite PLA natural, Polymaker), and the enclosure lid was made from laser cutting a 2mm-thick PMMA sheet (VLS 2.7, Universal Laser System).

Agar salt bridges were prepared mixing agarose (20-102, Apex Bioresearch) and DPBS (D8537, Sigma-Aldrich) at 4% w/v on a hot plate. Once fully melted, the molten agar was cast into a mold made from laser cutting a 3mm thick PMMA sheet on a glass slide. After agar was solidified, a safety razor was used to trim the excess agar, then the glass slide was removed to pop the molded salt bridges with a tweezer into a dish filled with PBS to keep agar bridges hydrated. Bridges with bubbles were discarded.

Ag|AgCl electrodes were prepared by cutting silver foil (AA11440GW, Fisher Scientific) into 8×23mm strips. Each electrode was plated in 1M KCl solution (P9541-1KG, Millipore Sigma) for 3 hours with a current of 3mA and a titanium wire cathode (00362-G1, Alfa Aesar). The coated portion should be around 8×15mm. Then the coated Ag|AgCl cathode and Ag anode were or inserted into the PCB lid or soldered to 24 awg wires (only for bi-axial stimulation). The electrodes had a charge capacity of 12mAh, no more than 8mAh of charge was used in each experiment to prevent faradaic reaction due to depletion of AgCl at the cathode.

### Joule heating measurement

Temperature rise due to joule heating was measured from placing a T-type thermocouple (5TC-KK-T-30-72, Omega) at the bottom of the well while running stimulation (See supplement fig). Various stimulation strengths up to twice the normal operating current were applied. The temperature rise readout (54 II B, Fluke) was less than 1C, see S4 for temperature deviation.

### pH stability measurement

Media from the stimulation 8 wells was measured after a stimulation delivery of 3V/cm for 3 hours using a portable pH meter (pH-33, Horiba). The stimulation well average was 7.29 (+/-0.12 std. div).

### Field uniformity verification through simulation and electrophoresis

Insert geometry was modeled in a finite element software (COMSOL 5.6) using material conductivity 1.6 S/m and relative permittivity of 80. It shows less than 10% variation in field a 6×6mm center, which aligns with many of similar devices’ design goals^16^. Actual field strength was measured by inserting two Ag|AgCl coated probes into the bottom of the insert. The measured voltage agrees with the COMSOL simulation. However, to further validate the result of the stimulation, we inject charged florescent beads into the stimulation well and measure its speed due to electrophoresis [SUPP]. Fluorescently labelled charged beads (F-8765, Thermo Fisher) was diluted 1:1000 in PBS and then added 1.5ml to the stimulation well. After device assembly as shown before, the setup was imaged using a confocal microscope (NL5, Confocal NL) with a 10x 0.3 NA objective in FITC channel with a frame rate of 5 frames/second.

Electrophoresis on charged fluorescent bead reach equilibrium with the stokes drag of the fluid. Therefore, beads speed is linearly proportional to the field strength. PIV analysis was then performed on the captured images.

### Cell culture

Primary mouse keratinocytes were provided by the Devenport Laboratory at Princeton University and cultured in E-medium (Nowak and Fuchs, 2009) supplemented with 15% serum (S11550, Atlanta Biologicals) and 50 μM calcium in the form of calcium chloride. Wild-type MDCK-II cells (courtesy of the Nelson Laboratory, Stanford University) were cultured in Dulbecco’s Modified Eagle’s Medium (D5523-10L, Sigma-Aldrich) with 1 g/L sodium bicarbonate, 10% fetal bovine serum (S11550, Atlanta Biologicals), and 1% penicillin–streptomycin (15140-122, Gibco). All cells were maintained at 37 °C under 5% CO2 and 95% relative humidity. Cells were split before reaching 70% of confluence for maintenance culture and were discarded after passage 30.

### Generation of TMEM154-eGFP MDCK Cellline

The generation and integration of the fluorescent TMEM154-eGFP fusion construct used in this study has been previously described^21^. In short, the construct was synthesized (GenScript) and cloned into a PiggyBAC transposon backbone, then stably integrated into MDCK-II cells via Lipofectamine LTX transfection (15338100, Thermo Fisher), followed by single-cell sorting (MA-900, Sony Biotechnology Inc.) and clonal expansion. Clones were screened for stable eGFP expression and proper membrane localization by flow cytometry (Invitrogen Attune NxT) and fluorescence microscopy.

### Tissue patterning and labeling

Tissue patterning was similar to previously published protocols^16,18,69^. 24 well plates (EW-01927-74, Cole-Parmer) were coated with Human Plasma Fibronectin Purified Protein (FC-010, Millipore Sigma) for keratinocyte experiment, or collagen IV (C5533-5MG, Sigma) for MDCK experiment. Each well was coated with 90μL of 50μg/ml of fibronectin or collagen under a 15mm circular glass coverslip (CLS-1760-015, Chemglass) for 30 minutes under 37C, then was washed three times with deionized water (DI). After allowing the dish to air dry, stencils were cut from 250-μm-thick silicone (HT-6240, Stockwell Elastomers) using a Silhouette Cameo vinyl cutter (Silhouette, USA) and transferred to the fibronectin-coated wells. 125μL of media was added around the edge of each well to keep it humidified during seeding and attachment.

Keratinocytes were concentrated at a concentration of 2.25 × 10^6^ cells/ml, and then pipetted into the stencil for ∼1000 cells/mm^2^, cares were taken not to scratch the fibronectin-coated substrate. Then the plate was incubated for 4 hours, allowing keratinocytes to attach to the substrate. After attachment, 500μL of media is added in each well, flooding the tissue. The stencils were lifted after 18 hours, which allows cells to form a confluent monolayer prior to stimulation. Cellbrite red (30023, Biotium) was added 5μL/ml of media, and each tissue is stained with 500μL of staining media for 30 minutes, then replaced with normal culture media for imaging. For nuclear tracking, Hoechst 33342 (D1306, Thermo Fisher) was used 1:1000 for 30 minutes. For actin labeling, cells were stained with SPY555-Actin (SC202, Spirochrome) 1:1000 for 30 minutes and were imaged with the probe in media.

MDCK cysts culture was described previously^13^. In short, MDCK cells were concentrated at a concentration of 2.25 × 10^5^ cells/ml in DMEM solution. Geltrex matrix (A1413201, Gibco) was mixed with the media at 1:1 ratio. 3.4μL of the resulting mixture was deposited onto the opening of the PDMS stencil. After solidifying for 30mins, 2.9μL of the mixture was deposited onto the first layer. The gel was then incubated for 1 hour before flooding with DMEM. The cysts were cultured for 96 hours and was imaged using Memglow 640 (MG04-10, Cytoskeleton. Inc). The sample was stained for 30 mins using 100nM concentration.

For transwell stimulations, approximately 86,000 TMEM154-eGFP MDCK is seeded to a 0.4μm pore size transwell membrane insert (353180, Corning). The samples were then incubated for 18 hrs before experiment. Samples were imaged in FITC channel.

### Device assembly

All components except salt bridges and inserts were sterilized by ethanol and then air dried in tissue culture hood. Agar salt bridges were sterilized by exposure to UV radiation for 5 minutes. The inserts were placed in sterile deionized water immediately after printing and were also UV sterilized. The 24 well plate is placed onto a hot plate (946C+, Soiiw) to keep cells at 37C during assembly and reduce focus drift due to thermal expansion in imaging. 2mL PBS was added for each well that did not contain tissues, and 1.5mL of media was added to each tissue well. Next, the inserts were carefully dropped into the wells. Then the plate was inspected from the bottom for any trapped bubbles, which may compromise field uniformity.

Next, agar salt bridges were inserted after drying the excess PBS onto a piece of Kimwipe (06-666 Kimberly-Clark Professional). The PCB lid was then dropped down with the electrodes aligned with each insert. For bi-axial stimulation, electrodes were inserted into the slits on the 3D printed insert and the wires were fixed using adhesive mounting putty (2600205, Loctite) for strain relief. The assembled platform was ready to transfer to the microscope while maintaining a sterile condition for the samples.

### Stimulation delivery and instrumentation

An 8-channel programmable current source was designed to minimize device footprint and provide easy stimulation control. It is capable of sourcing or sinking up to 20mA or 20V per channel. In short, each channel consists of a precision voltage-to-current transducer, analog switches for routing and to create (floating) negative voltages, as well as buffered ADCs for voltage and current samplings. The current command is generated through an 8-channel DAC with external voltage reference. Data logging and control were through serial communication between the computer and Teensy 4.0 MCU on the current source. The detailed block diagram is shown in Fig S1 and MATLAB GUI in Fig S2. Complete bill of materials, schematics, and fabrication files are on GitHub.

### Microscopy

All images were captured on an automated inverted microscope (Ti-2, Nikon) with motorized X-Y stage that was controlled using NIS-Element (Nikon Instruments). The microscope was incubated at 37C and 5% CO2 was perfused into the KENNEL enclosure. Fluorescence images were captured using a solid-state excitation source (Sola, Nikon) at 15% power and 4X/0.13 objective with an exposure of 400ms to a CMOS camera (Qi-2, Nikon). A 10x/0.45 objective was used for individual cell tracking. Images were taken at 5 minutes interval. For actin imaging, a 40X/1.4 oil objective was used on a 24-well glass button plate (P24-1.5H-N, CellVis). A spinning disk confocal (X-light V3, CrestOptics) was used in conjunction with the microscope to obtain images under high magnification. For cyst imaging, a 40X/1.25 silicone oil objective was used to provide extra working distance

### Image analysis and quantification

All images were processed using FIJI (https://imagej.net/software/fiji) and MATLAB (MathWorks) was used for data visualization and statistical analysis.

Particle image velocimetry (PIV) of the tissue was performed using PIVLab^70^, a MATLAB script for FFT-based PIV. Analysis was performed over the entirety of the tissue, excluding the edge, with an initial pass of 64×64 pixels and then 32×32 pixels with 50% overlap. The results were filtered using the script’s velocity-based validation. Vectors beyond 7 standard deviations were replaced by interpolated vector from adjacent vectors. The filtered results were imported to MATLAB for processing. Resulting migration speed and order data is smoothed by using a moving average filter with a window size of 5.

Tracking was performed for DAPI-labelled images using TrackMate^71^, an ImageJ plugin using simple LAP tracker. The results were filtered to exclude any incomplete track path and then were exported to a .XML file for MATLAB. A custom script was used to calculate the average migration track.

## Reference

1. Cortese, B., Palamà, I.E., D’Amone, S., and Gigli, G. (2014). Influence of electrotaxis on cell behaviour. Integr Biol 6, 817–830. 10.1039/c4ib00142g.

2. Zhao, M., Song, B., Pu, J., Wada, T., Reid, B., Tai, G., Wang, F., Guo, A., Walczysko, P., Gu, Y., et al. (2006). Electrical signals control wound healing through phosphatidylinositol-3-OH kinase-γ and PTEN. Nature 442, 457–460. 10.1038/nature04925.

3. Cohen, D.J., James Nelson, W., and Maharbiz, M.M. (2014). Galvanotactic control of collective cell migration in epithelial monolayers. Nature Mater 13, 409–417. 10.1038/nmat3891.

4. Allen, G.M., Mogilner, A., and Theriot, J.A. (2013). Electrophoresis of Cellular Membrane Components Creates the Directional Cue Guiding Keratocyte Galvanotaxis. Current Biology 23, 560–568. 10.1016/j.cub.2013.02.047.

5. Kennard, A.S., and Theriot, J.A. (2020). Osmolarity-independent electrical cues guide rapid response to injury in zebrafish epidermis. eLife 9, e62386. 10.7554/eLife.62386.

6. Zajdel, T.J., Shim, G., and Cohen, D.J. (2021). Come together: On-chip bioelectric wound closure. Biosensors and Bioelectronics 192, 113479. 10.1016/j.bios.2021.113479.

7. Song, J.W., Ryu, H., Bai, W., Xie, Z., Vázquez-Guardado, A., Nandoliya, K., Avila, R., Lee, G., Song, Z., Kim, J., et al. (2023). Bioresorbable, wireless, and battery-free system for electrotherapy and impedance sensing at wound sites. Science Advances 9, eade4687. 10.1126/sciadv.ade4687.

8. Sun, Y., Ferreira, F., Reid, B., Zhu, K., Ma, L., Young, B.M., Hagan, C.E., Tsolis, R.M., Mogilner, A., and Zhao, M. (2024). Gut epithelial electrical cues drive differential localization of enterobacteria. Nat Microbiol 9, 2653–2665. 10.1038/s41564-024-01778-8.

9. McCaig, C.D., Rajnicek, A.M., Song, B., and Zhao, M. (2005). Controlling cell behavior electrically: current views and future potential. Physiol Rev 85, 943–978. 10.1152/physrev.00020.2004.

10. Jiang, Y., Trotsyuk, A.A., Niu, S., Henn, D., Chen, K., Shih, C.-C., Larson, M.R., Mermin-Bunnell, A.M., Mittal, S., Lai, J.-C., et al. (2023). Wireless, closed-loop, smart bandage with integrated sensors and stimulators for advanced wound care and accelerated healing. Nat Biotechnol 41, 652–662. 10.1038/s41587-022-01528-3.

11. Sun, Y., Reid, B., Ferreira, F., Luxardi, G., Ma, L., Lokken, K.L., Zhu, K., Xu, G., Sun, Y., Ryzhuk, V., et al. (2019). Infection-generated electric field in gut epithelium drives bidirectional migration of macrophages. PLOS Biology 17, e3000044. 10.1371/journal.pbio.3000044.

12. Ferreira, F., Moreira, S., Zhao, M., and Barriga, E.H. (2025). Stretch-induced endogenous electric fields drive directed collective cell migration in vivo. Nat. Mater. 24, 462–470. 10.1038/s41563-024-02060-2.

13. Shim, G., Breinyn, I.B., Martínez-Calvo, A., Rao, S., and Cohen, D.J. (2024). Bioelectric stimulation controls tissue shape and size. Nat Commun 15, 2938. 10.1038/s41467-024-47079-w.

14. Mathews, J., and Levin, M. (2018). The body electric 2.0: recent advances in developmental bioelectricity for regenerative and synthetic bioengineering. Current Opinion in Biotechnology 52, 134–144. 10.1016/j.copbio.2018.03.008.

15. Gao, R., Zhao, S., Jiang, X., Sun, Y., Zhao, S., Gao, J., Borleis, J., Willard, S., Tang, M., Cai, H., et al. (2015). A large-scale screen reveals genes that mediate electrotaxis in Dictyostelium discoideum. Science Signaling 8, ra50–ra50. 10.1126/scisignal.aab0562.

16. Zajdel, T.J., Shim, G., Wang, L., Rossello-Martinez, A., and Cohen, D.J. (2020). SCHEEPDOG: Programming Electric Cues to Dynamically Herd Large-Scale Cell Migration. cels 10, 506–514.e3. 10.1016/j.cels.2020.05.009.

17. Leal, J., Shaner, S., Jedrusik, N., Savelyeva, A., and Asplund, M. (2023). Electrotaxis evokes directional separation of co-cultured keratinocytes and fibroblasts. Sci Rep 13, 11444. 10.1038/s41598-023-38664-y.

18. Shim, G., Devenport, D., and Cohen, D.J. (2021). Overriding native cell coordination enhances external programming of collective cell migration. Proc Natl Acad Sci U S A 118, e2101352118. 10.1073/pnas.2101352118.

19. Nwogbaga, I., Kim, A.H., and Camley, B.A. (2023). Physical limits on galvanotaxis. Phys. Rev. E 108, 064411. 10.1103/PhysRevE.108.064411.

20. Sarkar, A., Kobylkevich, B.M., Graham, D.M., and Messerli, M.A. (2019). Electromigration of cell surface macromolecules in DC electric fields during cell polarization and galvanotaxis. J. Theor. Biol. 478, 58–73.

21. Belliveau, N.M., Footer, M.J., Platenkamp, A., Rodriguez, C., Kim, H., Prinz, C.K., Loon, A.P. van, Lin, Y., Eustis, T.E., Chan, M.M., et al. (2025). Galvanin (TMEM154) is an electric-field sensor for directed cell migration. Preprint at bioRxiv, 10.1101/2024.09.23.614580 https://doi.org/10.1101/2024.09.23.614580.

22. Lyon, J.G., Carroll, S.L., Mokarram, N., and Bellamkonda, R.V. (2019). Electrotaxis of Glioblastoma and Medulloblastoma Spheroidal Aggregates. Sci Rep 9, 5309. 10.1038/s41598-019-41505-6.

23. Tai, G., Tai, M., and Zhao, M. (2018). Electrically stimulated cell migration and its contribution to wound healing. Burns & Trauma 6. 10.1186/s41038-018-0123-2.

24. Shaner, S., Savelyeva, A., Kvartuh, A., Jedrusik, N., Matter, L., Leal, J., and Asplund, M. Bioelectronic microfluidic wound healing: a platform for investigating direct current stimulation of injured cell collectives. Lab Chip 23, 1531–1546. 10.1039/d2lc01045c.

25. Matter, L., Abdullaeva, O.S., Shaner, S., Leal, J., and Asplund, M. (2024). Bioelectronic Direct Current Stimulation at the Transition Between Reversible and Irreversible Charge Transfer. Advanced Science 11, 2306244. 10.1002/advs.202306244.

26. Leal, J., Jedrusik, N., Shaner, S., Boehler, C., and Asplund, M. (2021). SIROF stabilized PEDOT/PSS allows biocompatible and reversible direct current stimulation capable of driving electrotaxis in cells. Biomaterials 275, 120949. 10.1016/j.biomaterials.2021.120949.

27. Sun, Y.-S. (2017). Studying Electrotaxis in Microfluidic Devices. Sensors (Basel) 17, 2048. 10.3390/s17092048.

28. Wu, J., and Lin, F. (2014). Recent Developments in Electrotaxis Assays. Adv Wound Care (New Rochelle) 3, 149–155. 10.1089/wound.2013.0453.

29. Pargar, F., Kolev, H., Koleva, D.A., and Breugel, K. van (2018). Microstructure, surface chemistry and electrochemical response of Ag|AgCl sensors in alkaline media. J. Mater. Sci. 53, 7527–7550. 10.1007/s10853-018-2083-0.

30. Schopf, A., Boehler, C., and Asplund, M. (2016). Analytical methods to determine electrochemical factors in electrotaxis setups and their implications for experimental design. Bioelectrochemistry 109, 41–48. 10.1016/j.bioelechem.2015.12.007.

31. Li, L., Li, G., Cao, Y., and Duan, Y.Y. (2021). A Novel Highly Durable Carbon/Silver/Silver Chloride Composite Electrode for High-Definition Transcranial Direct Current Stimulation. Nanomaterials-basel 11, 1962. 10.3390/nano11081962.

32. Zhao, S., Tseng, P., Grasman, J., Wang, Y., Li, W., Napier, B., Yavuz, B., Chen, Y., Howell, L., Rincon, J., et al. (2018). Programmable Hydrogel Ionic Circuits for Biologically Matched Electronic Interfaces. Adv. Mater. 30, 1800598. 10.1002/adma.201800598.

33. Driscoll, N., Erickson, B., Murphy, B.B., Richardson, A.G., Robbins, G., Apollo, N.V., Mentzelopoulos, G., Mathis, T., Hantanasirisakul, K., Bagga, P., et al. (2021). MXene-infused bioelectronic interfaces for multiscale electrophysiology and stimulation. Sci Transl Med 13, eabf8629. 10.1126/scitranslmed.abf8629.

34. Leal, J., Shaner, S., Matter, L., Böhler, C., and Asplund, M. (2023). Guide to Leveraging Conducting Polymers and Hydrogels for Direct Current Stimulation. Adv Mater Interfaces, 2202041. 10.1002/admi.202202041.

35. Wang, H., Zhao, Z., Liu, P., and Guo, X. (2022). Laser-Induced Graphene Based Flexible Electronic Devices. Biosensors 12, 55. 10.3390/bios12020055.

36. Dijk, G., Rutz, A.L., and Malliaras, G.G. (2020). Stability of PEDOT:PSS-Coated Gold Electrodes in Cell Culture Conditions. Adv Mater Technologies 5, 1900662. 10.1002/admt.201900662.

37. Boehler, C., and Asplund, M. (2018). PEDOT as a high charge injection material for low-frequency stimulation. 2018 40th Annu Int Conf Ieee Eng Medicine Biology Soc Embc 00, 2202–2205. 10.1109/embc.2018.8512597.

38. Cole, J., and Gagnon, Z. (2019). A flow-based microfluidic device for spatially quantifying intracellular calcium ion activity during cellular electrotaxis. Biomicrofluidics 13, 064107. 10.1063/1.5124846.

39. Sun, Y.-S., Peng, S.-W., and Cheng, J.-Y. (2012). In vitro electrical-stimulated wound-healing chip for studying electric field-assisted wound-healing process. Biomicrofluidics 6, 034117. 10.1063/1.4750486.

40. Peimani, A.R., Zoidl, G., and Rezai, P. (2018). A microfluidic device to study electrotaxis and dopaminergic system of zebrafish larvae. Biomicrofluidics 12, 014113. 10.1063/1.5016381.

41. Zhao, S., Zhu, K., Zhang, Y., Zhu, Z., Xu, Z., Zhao, M., and Pan, T. (2014). ElectroTaxis-on-a-Chip (ETC): an integrated quantitative high-throughput screening platform for electrical field-directed cell migration. Lab Chip 14, 4398–4405. 10.1039/C4LC00745J.

42. A Galvanotaxis Assay for Analysis of Neural Precursor Cell Migration Kinetics in an Externally Applied Direct Current Electric Field https://app.jove.com/v/4193/a-galvanotaxis-assay-for-analysis-neural-precursor-cell-migration.

43. Ralf, K., K, N.B., B, S.T., and Karen, E. (2020). Development of a multi-well-chip for studying 2D and 3D tumor cell migration and spheroid growth in electrical fields. Current Directions in Biomedical Engineering 6, 164–167. 10.1515/cdbme-2020-3042.

44. Electric Field-controlled Directed Migration of Neural Progenitor Cells in 2D and 3D Environments https://app.jove.com/v/3453/electric-field-controlled-directed-migration-neural-progenitor-cells.

45. Snyder, S., DeJulius, C., and Willits, R.K. (2017). Electrical Stimulation Increases Random Migration of Human Dermal Fibroblasts. Ann Biomed Eng 45, 2049–2060. 10.1007/s10439-017-1849-x.

46. Huang, C.-W., Cheng, J.-Y., Yen, M.-H., and Young, T.-H. (2009). Electrotaxis of lung cancer cells in a multiple-electric-field chip. Biosensors and Bioelectronics 24, 3510– 3516. 10.1016/j.bios.2009.05.001.

47. Zhang, Y., Lee, R.M., Zhu, Z., Sun, Y., Zhu, K., Xu, Z., Lin, F., Pan, T., Losert, W., and Zhao, M. (2023). Protocol for electrotaxis of large epithelial cell sheets. STAR Protocols 4, 102288. 10.1016/j.xpro.2023.102288.

48. Microfluidic-based Electrotaxis for On-demand Quantitative Analysis of Caenorhabditis elegans’ Locomotion https://app.jove.com/v/50226/microfluidic-based-electrotaxis-for-on-demand-quantitative-analysis.

49. Utilizing Custom-designed Galvanotaxis Chambers to Study Directional Migration of Prostate Cells https://app.jove.com/v/51973/utilizing-custom-designed-galvanotaxis-chambers-to-study-directional.

50. Tai, G., Reid, B., Cao, L., and Zhao, M. (2009). Electrotaxis and Wound Healing: Experimental Methods to Study Electric Fields as a Directional Signal for Cell Migration. In Chemotaxis: Methods and Protocols, T. Jin and D. Hereld, eds. (Humana Press), pp. 77–97. 10.1007/978-1-60761-198-1_5.

51. Zhang, Y., Xu, G., Wu, J., Lee, R.M., Zhu, Z., Sun, Y., Zhu, K., Losert, W., Liao, S., Zhang, G., et al. (2022). Propagation dynamics of electrotactic motility in large epithelial cell sheets. iScience 25. 10.1016/j.isci.2022.105136.

52. Tran, V., Zhang, X., Cao, L., Li, H., Lee, B., So, M., Sun, Y., Chen, W., and Zhao, M. (2013). Synchronization Modulation Increases Transepithelial Potentials in MDCK Monolayers through Na/K Pumps. PLOS ONE 8, e61509. 10.1371/journal.pone.0061509.

53. Abasi, S., Jain, A., Cooke, J.P., and Guiseppi-Elie, A. (2023). Electrically stimulated gene expression under exogenously applied electric fields. Front. Mol. Biosci. 10. 10.3389/fmolb.2023.1161191.

54. Kramer, N., Walzl, A., Unger, C., Rosner, M., Krupitza, G., Hengstschläger, M., and Dolznig, H. (2013). In vitro cell migration and invasion assays. Mutat Res 752, 10–24. 10.1016/j.mrrev.2012.08.001.

55. Hulkower, K.I., and Herber, R.L. (2011). Cell Migration and Invasion Assays as Tools for Drug Discovery. Pharmaceutics 3, 107–124. 10.3390/pharmaceutics3010107.

56. Saw, T.B., Gao, X., Li, M., He, J., Le, A.P., Marsh, S., Lin, K., Ludwig, A., Prost, J., and Lim, C.T. (2022). Transepithelial potential difference governs epithelial homeostasis by electromechanics. Nat. Phys. 18, 1122–1128. 10.1038/s41567-022-01657-1.

57. Zhang, Y., Xu, G., Lee, R.M., Zhu, Z., Wu, J., Liao, S., Zhang, G., Sun, Y., Mogilner, A., Losert, W., et al. (2017). Collective cell migration has distinct directionality and speed dynamics. Cell Mol Life Sci 74, 3841–3850. 10.1007/s00018-017-2553-6.

58. Reid, B., Song, B., McCaig, C.D., and Zhao, M. (2005). Wound healing in rat cornea: the role of electric currents. The FASEB Journal 19, 379–386. 10.1096/fj.04-2325com.

59. Arnold, C.E., Rajnicek, A.M., Hoare, J.I., Pokharel, S.M., Mccaig, C.D., Barker, R.N., and Wilson, H.M. (2019). Physiological strength electric fields modulate human T cell activation and polarisation. Sci Rep 9, 17604. 10.1038/s41598-019-53898-5.

60. Lin, F., Baldessari, F., Gyenge, C.C., Sato, T., Chambers, R.D., Santiago, J.G., and Butcher, E.C. (2008). Lymphocyte Electrotaxis in vitro and in vivo. J Immunol 181, 2465– 2471. 10.4049/jimmunol.181.4.2465.

61. Hinkle, L., McCaig, C.D., and Robinson, K.R. (1981). The direction of growth of differentiating neurones and myoblasts from frog embryos in an applied electric field. The Journal of Physiology 314, 121–135. 10.1113/jphysiol.1981.sp013695.

62. Levin, M. (2021). Bioelectric signaling: Reprogrammable circuits underlying embryogenesis, regeneration, and cancer. Cell 184, 1971–1989. 10.1016/j.cell.2021.02.034.

63. Taeger, C.D., Friedrich, O., Dragu, A., Weigand, A., Hobe, F., Drechsler, C., Geppert, C.I., Arkudas, A., Münch, F., Buchholz, R., et al. (2015). Assessing viability of extracorporeal preserved muscle transplants using external field stimulation: a novel tool to improve methods prolonging bridge-to-transplantation time. Sci Rep 5, 11956. 10.1038/srep11956.

64. Naranjo-Hernández, D., Reina-Tosina, J., Roa, L.M., Barbarov-Rostán, G., Calvillo-Arbizu, J., Talaminos-Barroso, A., Pérez-Valdivia, M.Á., and Medina-López, R.A. (2025). Development and Validation in Porcine and Human Models of a Bioimpedance Spectroscopy System for the Objective Assessment of Kidney Graft Viability. Sensors 25, 2871. 10.3390/s25092871.

65. Berkelmann, L., Bader, A., Meshksar, S., Dierks, A., Hatipoglu Majernik, G., Krauss, J.K., Schwabe, K., Manteuffel, D., and Ngezahayo, A. (2019). Tumour-treating fields (TTFields): Investigations on the mechanism of action by electromagnetic exposure of cells in telophase/cytokinesis. Sci Rep 9, 7362. 10.1038/s41598-019-43621-9.

66. Wenger, C., Miranda, P.C., Salvador, R., Thielscher, A., Bomzon, Z., Giladi, M., Mrugala, M.M., and Korshoej, A.R. (2018). A Review on Tumor-Treating Fields (TTFields): Clinical Implications Inferred From Computational Modeling. Ieee Rev Biomed Eng 11, 195–207. 10.1109/rbme.2017.2765282.

67. Hottinger, A.F., Pacheco, P., and Stupp, R. (2016). Tumor treating fields: a novel treatment modality and its use in brain tumors. Neuro-oncology 18, 1338–1349. 10.1093/neuonc/now182.

68. Hong, P., Kudulaiti, N., Wu, S., Nie, J., and Zhuang, D. (2021). Tumor treating fields: a comprehensive overview of the underlying molecular mechanism. Expert Rev Mol Diagn 22, 19–28. 10.1080/14737159.2022.2017283.

69. Heinrich, M.A., Alert, R., Wolf, A.E., Košmrlj, A., and Cohen, D.J. (2022). Self-assembly of tessellated tissue sheets by expansion and collision. Nat Commun 13, 4026. 10.1038/s41467-022-31459-1.

70. (PDF) Particle Image Velocimetry for MATLAB: Accuracy and enhanced algorithms in PIVlab (2024). ResearchGate. 10.5334/jors.334.

71. Ershov, D., Phan, M.-S., Pylvänäinen, J.W., Rigaud, S.U., Le Blanc, L., Charles-Orszag, A., Conway, J.R.W., Laine, R.F., Roy, N.H., Bonazzi, D., et al. (2022). TrackMate 7: integrating state-of-the-art segmentation algorithms into tracking pipelines. Nat Methods 19, 829–832. 10.1038/s41592-022-01507-1.

