## Supplement figures for "SCHEPHERD: A universal platform for high-throughput, high-resolution, and programmable control of cell behavior through bioelectric stimulation"

Supplemental figures

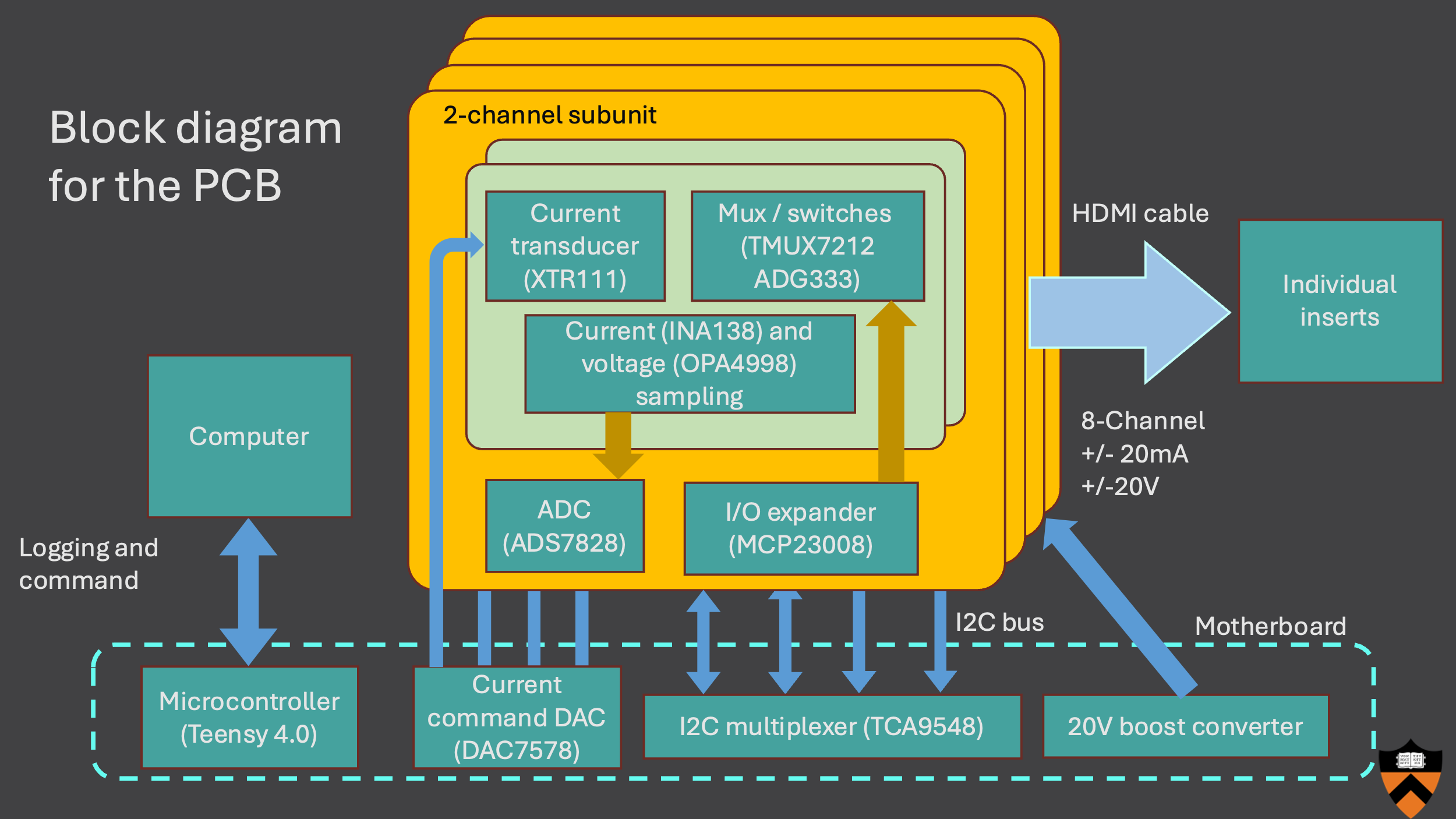

**Fig. S1. PCB block diagram**
The multi-channel programmable current source uses a motherboard that generates the current command signal through a precision DAC. The mother board can host up to 4 subunit modules, with multiplexed I2C communication to avoid address conflict. All command is relayed through a microcontroller that connects to a PC via serial. Each subunit consists of two independent current sources and voltage/sampling circuit through I2C. Polarity is determined by analog multiplexers and switches, which are also controlled by I2C. Different subunits may be used to provide higher resolution at lower current or to achieve higher current output. All 8 channels are connected to the electrode PCB via a HDMI cable, minimizing assembly time and streamlining cable management.

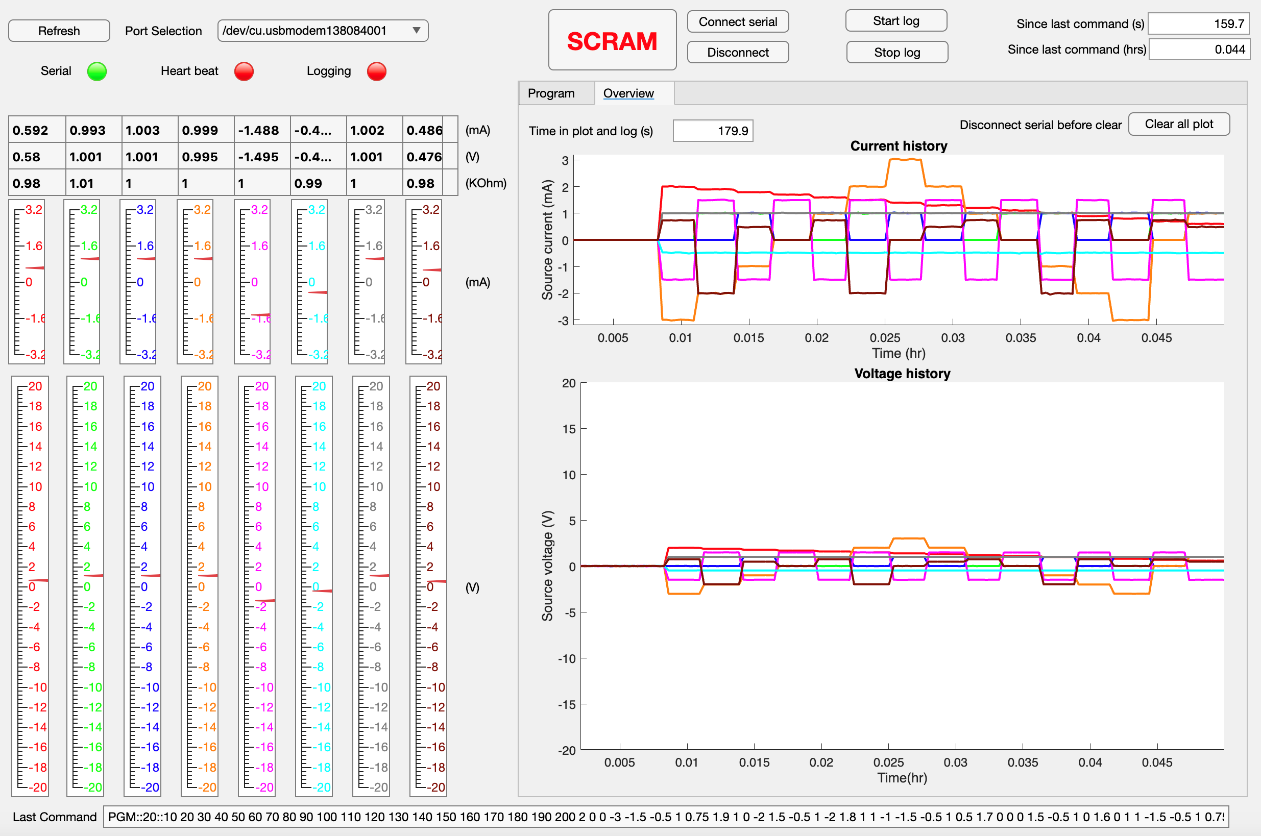

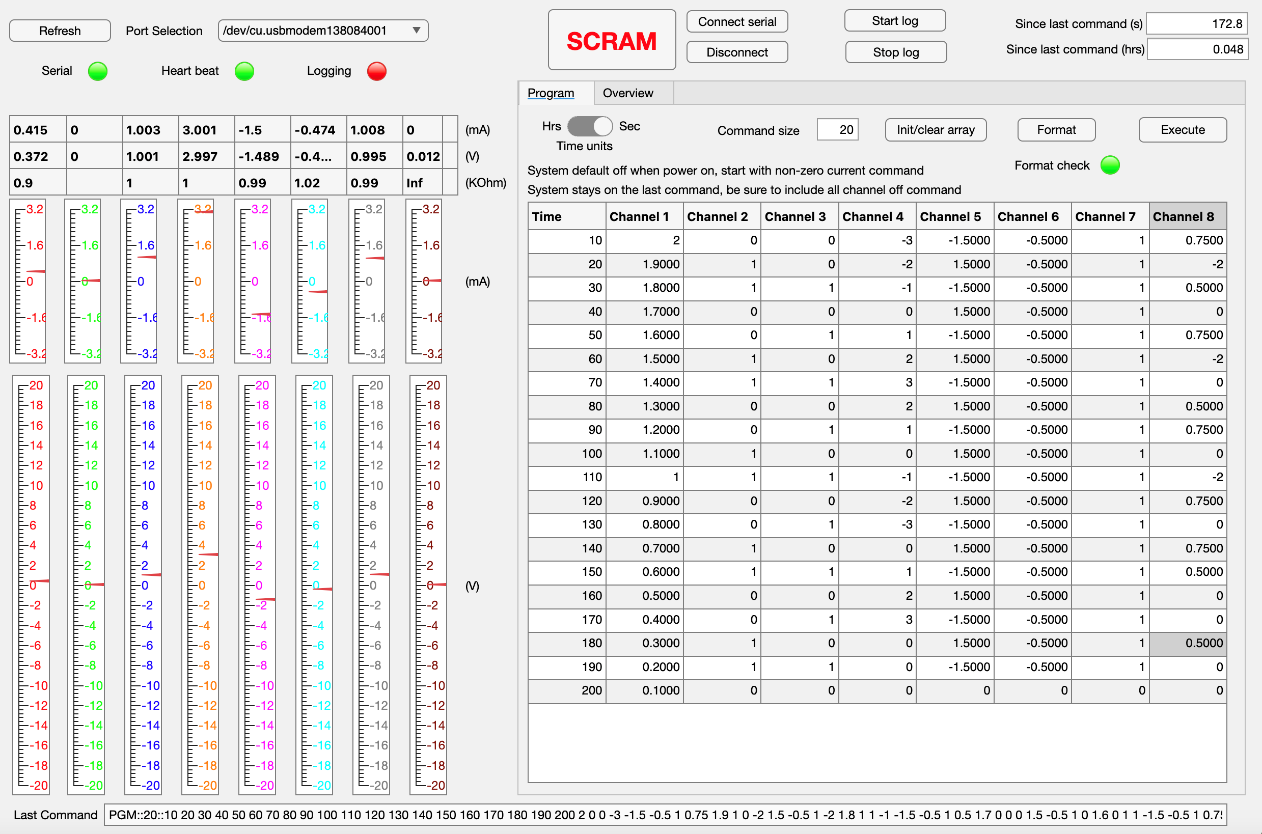

**Fig. S2. Multi-channel current source programming interface for complex stimulation control**
This MATLAB-based app provides flexibility for programming the current source. The top image is a screenshot of current and voltage readbacks on 8 channels during stimulation, while the bottom image shows programming interface with variable command length and current polarity. Linear gauges on the left side provide ease of comparing across channels.

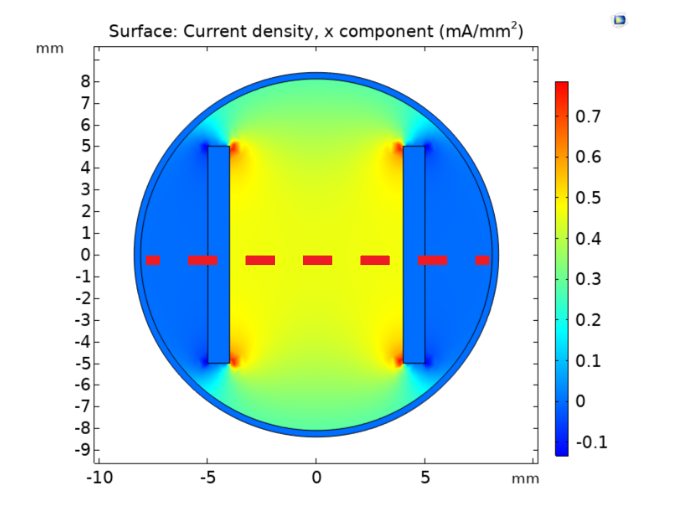

**Fig. S3. COMSOL simulation for flat insert**

COMSOL simulation for the flat insert with 1.5mA of stimulation current, showing a uniform stimulation zone.

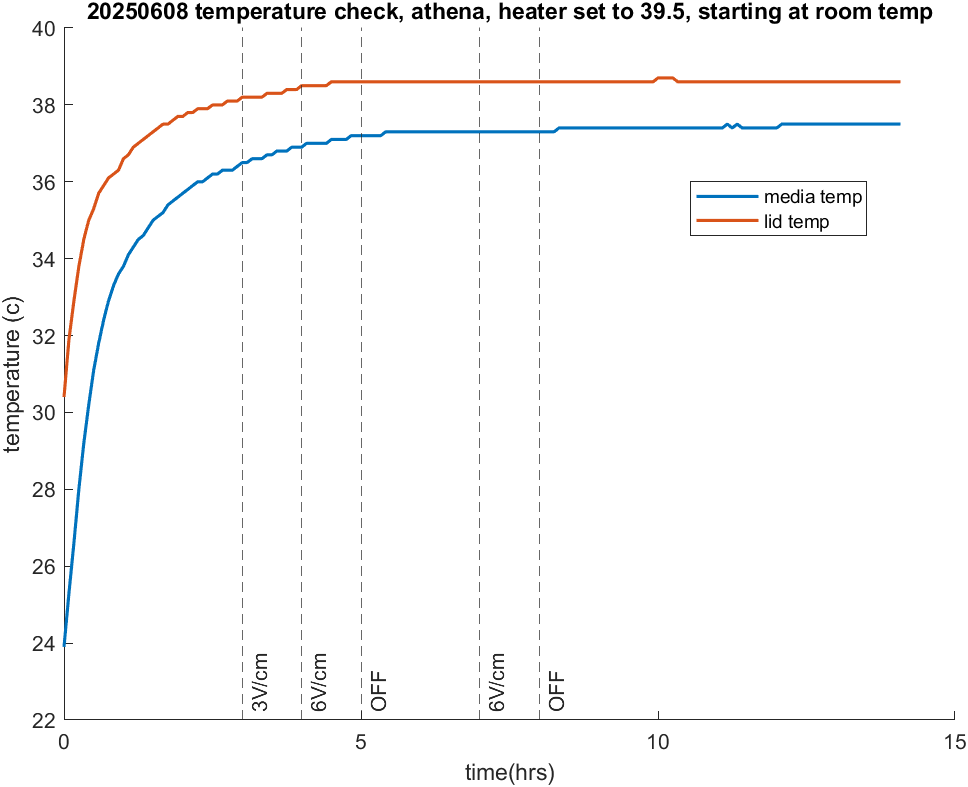

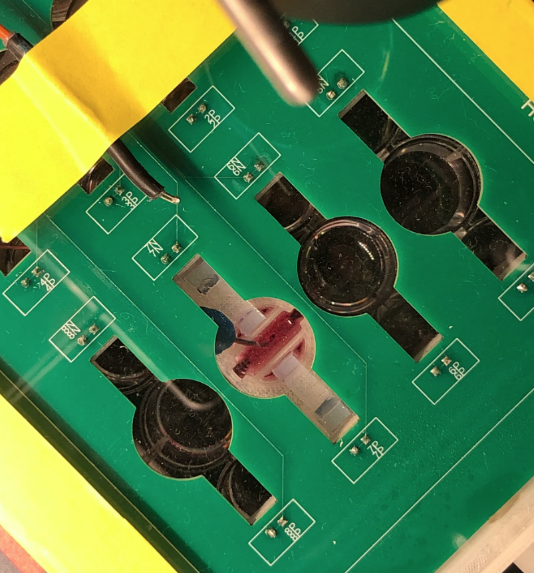

**S4. Temperature stability and joule heating validation**

To ensure no significant joule heating occurs during stimulation, two temperature probes are placed on the top of the acrylic lid (yellow box) and inside the stimulation insert with media (blue box). Different stimulation strength is annotated on the left plot. The results show no significant joule heating (less than 0.5C) after the insert was equilibrated from room temperature.

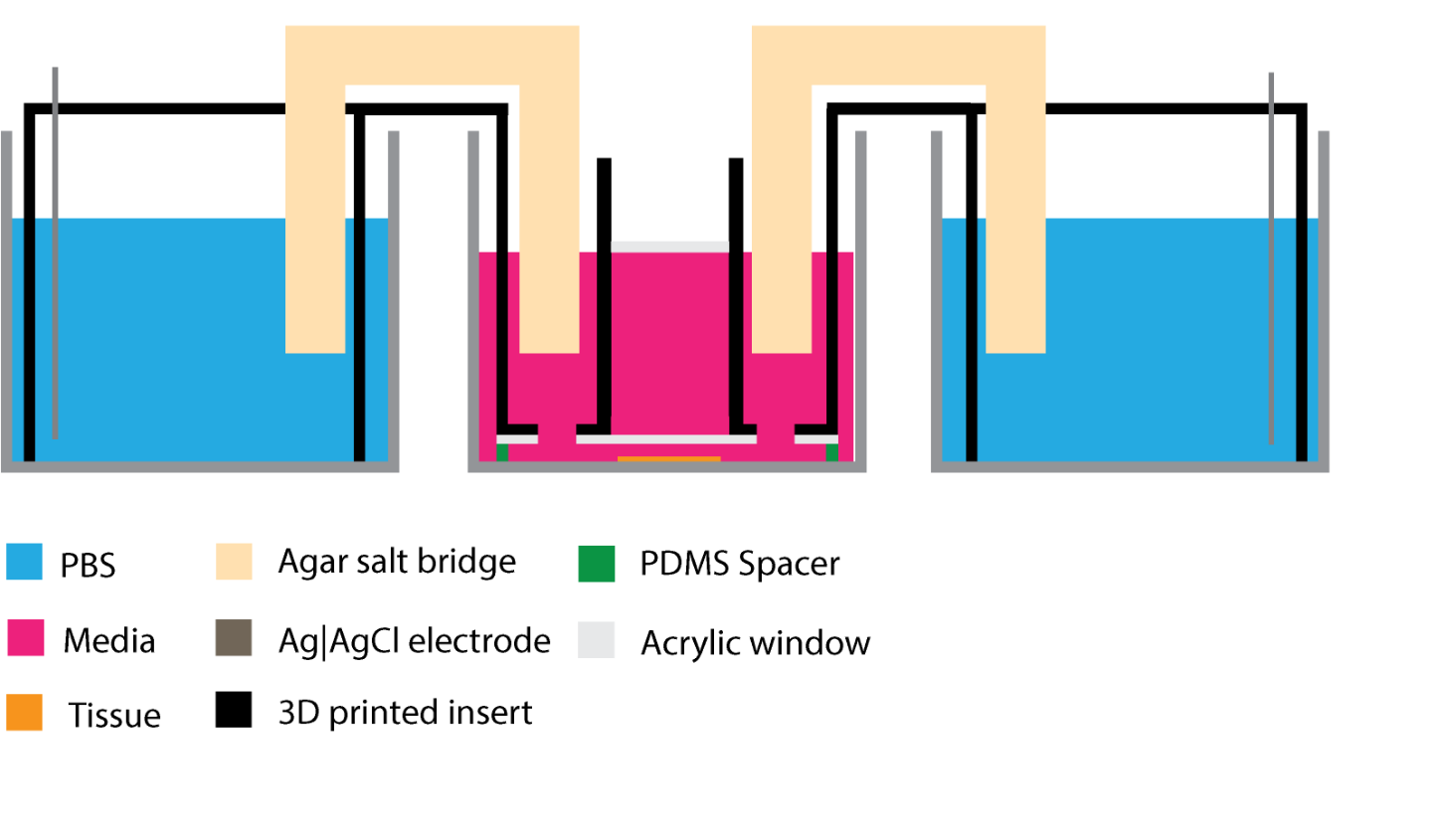

**Fig. S5. Transmitted light insert schematic with transparent inner window**

The SCHEPHERD system can be modified for transmitted light imaging. Two laser-cut acrylic blocks are attached to the insert (highlighted by the green arrows) for transmitted light path. The top acrylic block also flats out the meniscus caused by the surface tension. In this case, 12 well plates are used to provide larger imaging windows, which allows up to 4 samples stimulated. 3.6 mA current will generate 1.5 V/cm field strength across a 5x5 mm area.

**Supplemental Video Captions**

**Video S1 – Uniform uniaxial stimulation**
A 1.5 × 6 mm mouse keratinocyte tissue undergoing electrotaxis at 3 V/cm.

**Video S2 – Uniaxial stimulation with transmitted light imaging**
A 1.5 × 6 mm MDCK tissue undergoing electrotaxis at 3 V/cm with phase contrast imaging.

**Video S3 – 3D Cyst electrotaxis and electro-inflation**
An MDCK cyst undergoing electrotaxis and electro-inflation within a hydrogel.

**Video S4 – Colony edge dynamics under paracellular stimulation**
An MDCK colony exhibiting expansion and retraction with paracellular stimulation in a transwell device.

**Video S5 – Large MDCK colony under paracellular stimulation**
Paracellular stimulation of MDCK but with a larger colony.

**Video S6 – Amplitude sweep with MDCK replicates**
Three 1 × 6 mm MDCK tissues, spaced 1 mm apart, used for amplitude sweep experiments.

**Video S7 – Bi-axial tissue-level simulation**
A mouse keratinocyte tissue responding to a series of 45° and 90° field commands, producing a triangular migration trajectory over 8 hours.

**Video S8 – Cytoskeletal rearrangement under bi-axial stimulation**
A single mouse keratinocyte stained with F-actin responding to sequential perpendicular field commands, showcasing dynamic cytoskeletal reorganization.

**Video S9 – Complex dynamics and tissue sculpting in quadrupolar fields**
A 5 × 5 mm differentiated mouse keratinocyte tissue exposed to a quadrupolar field insert, resulting in complex migration dynamics and local density changes.

**Video S10 – Localized electrotaxis using modified ceiling**
An MDCK tissue responding to localized field delivery, where only the “zig-zag” portion migrates.
